# Genomic rearrangement of the capsule operon in a phylogenetically distinct cluster of the multi-drug resistant *Escherichia coli* lineage ST131

**DOI:** 10.64898/2025.12.15.689991

**Authors:** Isabelle Potterill, Rebecca J. Hall, Robert A. Moran, Elizabeth A. Cummins, Alan McNally

**Affiliations:** Institute of Microbiology and Infection, School of Infection, Inflammation, and Immunology, College of Medicine and Health, University of Birmingham; School of Life and Environmental Sciences, Faculty of Science, University of Sydney; Ineos Oxford Institute, Department of Biology, Life and Mind Building, University of Oxford

## Abstract

*Escherichia coli* ST131 is a dominant cause of multi-drug resistant extra-intestinal infections in humans. The lineage has a three-clade phylogenetic structure of which clade C has gained resistance to multiple classes of antibiotics and is globally disseminated. As well as antimicrobial resistance, a number of virulence factors contribute to the success of ST131 clade C, including the production of capsule. Previous work on a limited number of strains has shown that the capsule operon in ST131 is highly variable. In this study we analysed the capsule genetics of *E. coli* ST131 by interrogating 3,033 genomes. Our data show a clonal distribution of capsule types and identify a monophyletic group of ST131 clade C strains which are characterised by an atypical capsule operon with highly divergent alleles of the key *kpsM* and *kpsT* genes. Our analysis suggests there is ongoing evolution within clade C of ST131 which looks to be driven not by continuing evolution of antimicrobial resistance, but by host-pathogen interactions.

## Introduction

Capsule is an important virulence factor in pathogenic *Escherichia coli*, improving bacterial persistence in the environment, and providing advantages in host colonisation such as immune evasion (1). The capsule layer is formed of long chain polysaccharides that are moved across the membrane by large membrane spanning translocation complexes (2, 3). The similarity of some capsule polysaccharides to human glycoproteins helps bacteria to evade the immune system, preventing full activation of innate host defences and suppressing phagocytosis (4). The highly hydrated nature of the capsular layer protects the bacteria from environmental stresses such as desiccation and antimicrobials, as well as allowing for adaptation to new environments. Capsule can also affect DNA exchange, phage predation, efficiency of antimicrobial peptides, and plasmid uptake (5-7).

Over 80 serologically unique capsule types have been identified in *E. coli* (8). They are categorised into four groups according to biochemical and genetic properties. These groups follow two distinct transport pathways. Groups 1 and 4 are *wzy*-dependent, where the polysaccharide repeating unit is built in the cytoplasm before being transported to the inner membrane by a Wzx flippase. In the periplasm it is polymerised by Wzy before transportation across the outer membrane (OM) (9). Groups 2 and 3 are ATP-binding cassette (ABC) transporter dependent. The entire polysaccharide is assembled on a lipid carrier before being transported across the inner membrane by an ABC transporter (1) and then translocated across the OM in the same manner as the *wzy*-dependent pathway (9). Identification of capsule types involves using a panel of antisera (10), however, this is labour intensive and no longer commercially available as a service. There have been recent developments in *in silico* methods to genotype capsules in *E. coli* allowing for quicker identification and evaluation (6, 7). This has also led to the identification of novel and hitherto uncharacterised capsule types, adding further complexity to the capsule landscape of *E. coli*. Despite this complexity, there are clear associations between pathotypes and capsule types, with group 2 capsule types specifically associated with extra-intestinal pathogenic *E. coli* (ExPEC) (11, 12).

The operon of group 2 capsules comprises a three-part structure (13) - Figure 1. Regions 1 and 3 are conserved across group 2 and contain capsular polysaccharide genes that make up the transport complex. Region 1 includes genes *kpsFEDUCS* and region 3 *kpsM* and *kpsT*. Region 2 is the k-antigen determining region unique to the capsule polysaccharide locus (1). Region 1 genes *kpsD* and *kpsE* encode proteins to translocate the polysaccharide: *kpsD* encoding an outer membrane polysaccharide export protein and *kpsE* an inner membrane polysaccharide copolymerase protein (10). *kpsS* and *kpsC*, encode β-Kdo transferases polymerizing the oligosaccharides. *kpsS* transfers the first β-Kdo residue with the remaining transferred by *kpsC* (14). *kpsF* and *kpsU* encode enzymes for production of the CMP-Kdo. *kpsF* encodes a d-arabinose 5-phosphate isomerase, which converts d-ribulose 5-phosphate to d-arabinose 5-phosphate. *kpsU* is a CMP-Kdo synthetase (10). The *kpsMT* pair in region 3 facilitates the transportation of polysaccharides across the inner membrane. The KpsM protein is the integral inner membrane component and the KpsT protein an ATPase (15). In the normal arrangement of the operon, region 1 is transcribed from a promoter (PR1) 225bp upstream of *kpsF* and region 3 is transcribed from a promoter (PR3) 741bp upstream from *kpsM* (16, 17). These promoters are both temperature sensitive (17). The untranslated region between the promoter and *kpsM* is also believed to contribute to this temperature regulation (16). Transcription of region 2 genes is RfaH mediated with an operon polarity suppressor (ops) site found upstream of *kpsM* (18). RfaH is an anti-terminator allowing the transcription to continue beyond region 3, and therefore the expression of the k-antigen defining genes.

**Figure 1.**
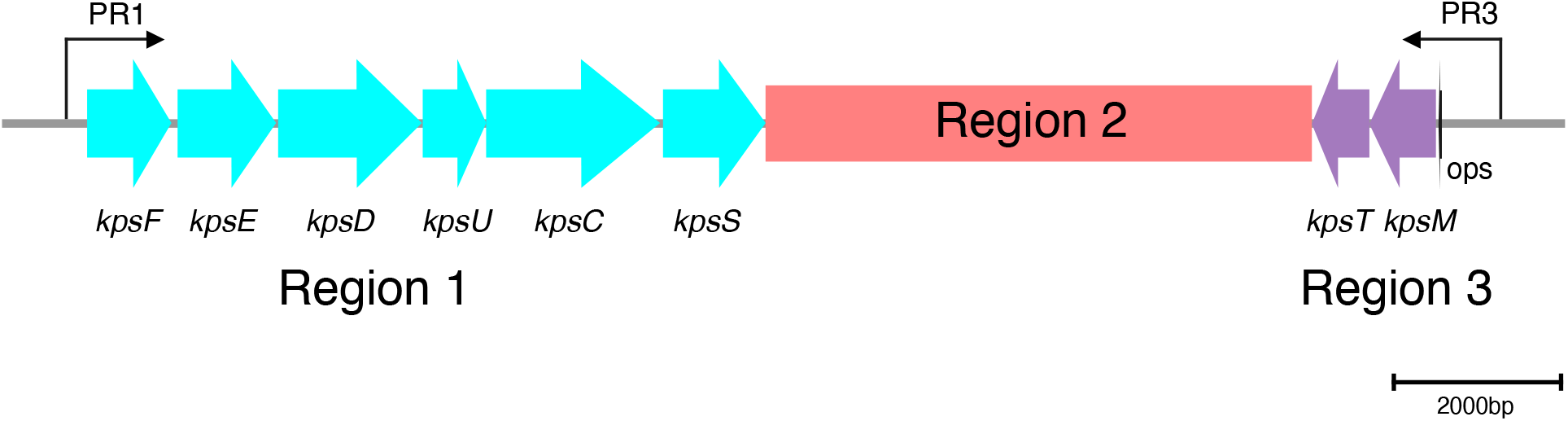
The conventional genetic structure of the group 2 *kps* operon. Three regions are represented with conserved region 1 and 3 flanking serotype-specific region 2. The genes within region 2 are unique to each k-antigen. PR1 represents the promoter upstream of *kpsF* which transcribes region 1. PR3 represents the promoter upstream of *kpsM* which transcribes region 3. Transcription of region 2 genes is RfaH mediated with an operon polarity suppressor (ops).

Previous work by our group, looking at a small number of genomes, identified surprising levels of diversity of the capsule regions of the pandemic multi-drug resistant (MDR) ExPEC lineage sequence type (ST) 131 (19). This diversity was surprising given that the MDR clade of the ST131 lineage (clade C) has relatively recently emerged and was considered largely clonal (20). A more recent paper has been published looking at a large collection of ExPEC genomes, showing significant variation in capsule genetics across the ExPEC lineages (6). Within this dataset we noticed a specific subset of the ST131 lineage which appeared to show a switch in capsule type. Capsule typing is calculated predominately on the variation of genes in region 2, the k-antigen defining region (21). By looking at a collection of 3,033 ST131 genomes we undertook an in-depth characterisation of the diversity of the capsule region across ST131, and specifically characterised a capsule switch in a phylogenetically distinct cluster within *E. coli* ST131 clade C. We show that this switch is characterised by a divergent *kpsM* variant that forms part of an atypical capsule operon.

## Methods

### Capsule locus typing of a curated genome collection of *E. coli* ST131

A collection of 20,577 *E. coli* genomes downloaded from Enterobase (22) was used in this study. The genome collection has previously been characterised by Cummins et al. 2019 (23). The collection consists of 21 STs from 7 phylogroups aiming to cover the diversity of the species. This work focusses on the 3,033 ST131 genomes within the collection (supplementary dataset).

Using the downloaded assembled ST131 genome files, the number of *kpsM* and *kpsT* alleles across ST131 was calculated using BLASTn (v2.5.0+) (24, 25). All nucleotide variants were translated into amino acids using The Sequence Manipulation Suite v2 (26). Both the nucleotide and the amino acid sequences were aligned using MAFFT online (v7.526) using default parameters (27). Alignments were visualised with Jalview (v2.11.4.1) (28) to look at the distribution of the differences across the alleles at both a nucleotide and an amino acid level. Single nucleotide polymorphism (SNP) distances between variants were calculated using snp-dists (v0.8.2) (https://github.com/tseemann/snp-dists) with default parameters on the nucleotide alignments. Comparison of the nucleotide identity between the capsule region of the EC958 (29) reference genome and divergent alleles was calculated with a global alignment tool (EBI – EMBOSS needle) (30). Online BLASTn searches were carried out, first with default parameters and secondly, excluding *E. coli*, to assess if there was evidence of the divergent *kpsM* allele being present in other species. The sequence type and phylogroup of *E. coli* matches were determined with mlst (v2.23.0) (https://github.com/tseemann/mlst) (31) and Clermontyping (v24.02) (32) respectively with default parameters. The entire collection of *E. coli* consisting of 21 STs was screened for all *kpsM* variants using BLASTn. Capsule typing was performed on assemblies from the entire collection with Kaptive (v3.0.0b6) (33) using the *E. coli* Group 2 and Group 3 capsular K-typing database (v3.0.0) (6).

### Phylogenetic reconstruction of capsule gene distribution

To understand the phylogenetic distribution of *kpsM* and *kpsT* variants across the lineage, a phylogenetic tree of the 3,033 ST131 genomes was built. A core gene alignment was created using panaroo (34) and raxmlHPC-PTHREADS-AVX (v8.2.12) (35) run with the GTRGAMMA model. SNP distances were calculated between *kpsM* and *kpsT* as above. All *kpsM* and *kpsT* variants were then manually clustered according to SNP distance for visualisation. Initially, clusters contained variants where all members were within ten SNPs of each other. A small number of variants were not assigned an initial cluster. These were manually added to a cluster if they were fewer than 11 SNPs away from at least one member of that cluster. These clusters, as well as capsule types, were mapped to the phylogenetic tree and visualised in iTOL (36). Concordance between *kpsM* and *kpsT* alleles to each other and to KL type was calculated using an adjusted rand index using R package mcclust (37) in R (v4.5.0).

### Long-read sequencing of divergent capsule type strains

Five bacterial strains were used from our strain collection, three were located within the identified clone containing the divergent capsule type (MVAST198 (38), MVAST348 (38), MVAST350 (38)), and two from outside the clone but within ST131 (USVAST219 (39), USVAST065 (39)). Single colonies of these five bacterial strains were taken, following overnight growth on LB agar, and added to 3 mL LB broth (Sigma-Aldrich, United Kingdom). These were grown overnight in a shaking incubator at 37 °C with 180 rpm. DNA extraction was performed using the Monarch Genomic DNA Purification kit (New England Biolabs) and quantified using the Qubit Broad Range dsDNA kit (ThermoFisher). The SQK-NBD114 ligation barcoding kit was used and genomic DNA sequenced on R10.4.1 flow cells using the GridION (ONT), with the addition of bovine serum albumin as recommended by the manufacturer. Reads were assembled using hybracter (40). Assemblies were annotated with Bakta (41) and visualised with Artemis (42). Comparisons between genomes were conducted using BLAST and Artemis.

### RNA sequencing of novel capsule type strains

Biological triplicates of four bacterial strains (MVAST198, MVAST348, MVAST350 and USVAST219) were submitted for RNA sequencing. Four isolates were streak plated on LB agar and grown at 37°C overnight. Single colonies were taken and added to 3 mL LB broth (Sigma-Aldrich, United Kingdom) and grown overnight in a shaking incubator at 37°C with 180 rpm. 250 µL of culture was aliquoted into 25 mL of fresh LB media and incubated at 37°C with 180 rpm in the shaking incubator until mid-log with an optical density at OD600 of approximately 0.5. 5mL of the sample was then centrifuged for 10 minutes at 4000 rpm, the supernatant poured off, and 1mL of PBS added. The samples were centrifuged again for 5 minutes at 13,200 rpm. The supernatant was removed, and the pellet was snap frozen. The frozen cell pellets were sent to GENEWIZ from Azenta Life Sciences (Frankfurt, Germany) using their standard RNA sequencing service.

Kallisto (v0.46.0) (https://github.com/pachterlab/kallisto) was used to obtain read counts for each gene, referenced against long-read assemblies assembled as above. Annotations were prepared with Bakta and converted from Genbank to kallisto with genbank_to_kallisto.py (https://github.com/AnnaSyme/genbank_to_kallisto.py). Read counts were normalised in each sample with respect to read counts in housekeeping gene *adk*.

## Results

### Capsule typing of a collection of *E. coli* ST131 genomes identifies a phylogenetically distinct cluster with a unique capsule type

Kaptive was run on our curated collection of 3,033 ST131 genomes to identify the *in silico* capsule type. Out of 3,033 isolates, 3,030 had capsule operons detected of which 2,914 were typeable and the remaining 105 genomes had untypeable capsule operons. For a given K-locus the genotype is reported with a KL-type (KL) and the inferred phenotype with a K-type (K). The K-type and KL-type, alongside *kpsM* and *kpsT* variants, were mapped to the phylogenetic tree of our ST131 genome collection (Fig 2).

**Figure 2.**
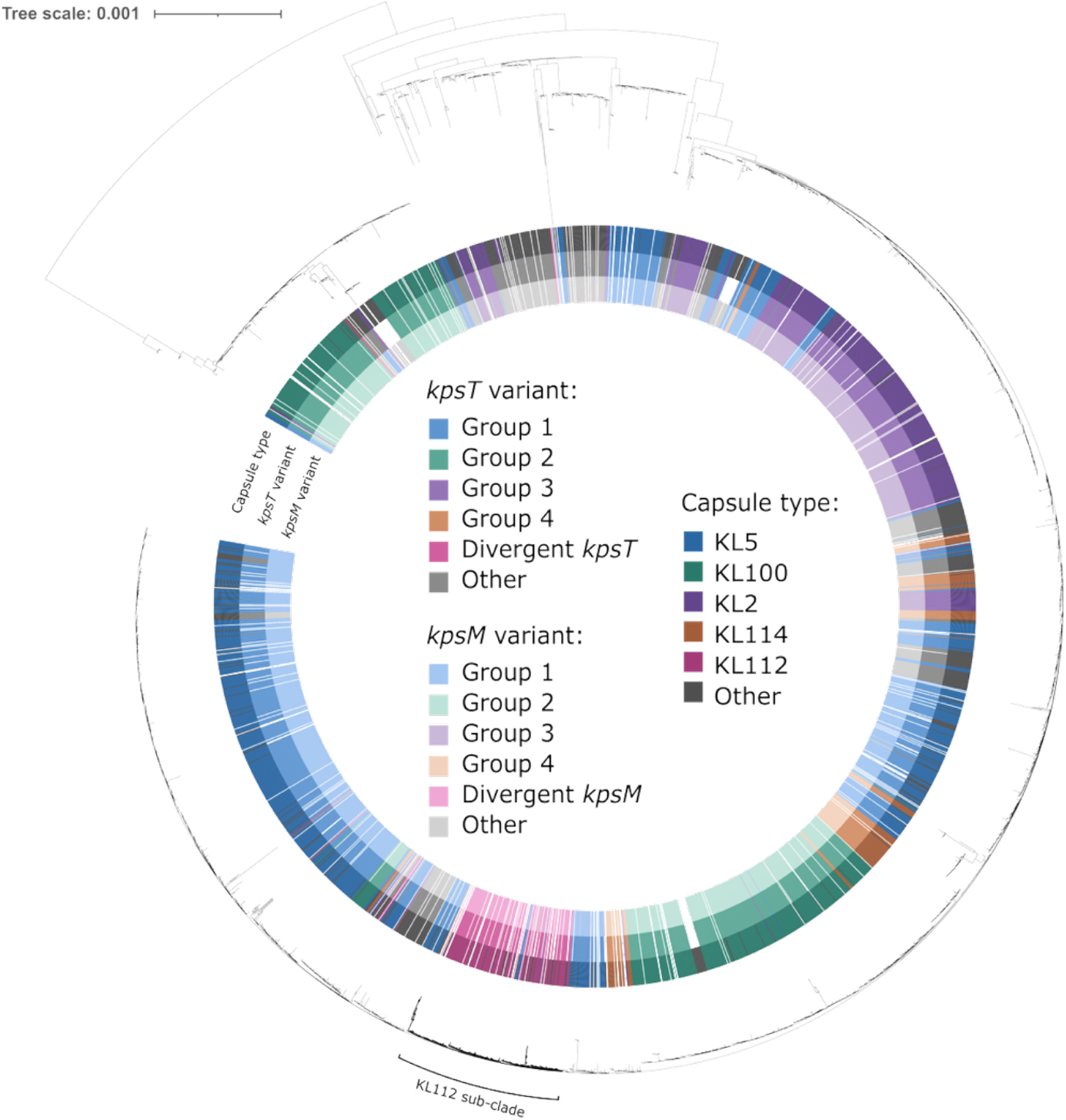
A phylogenetic tree of 3,033 ST131 *Escherichia coli* isolates. The outer ring represents the capsule type (as determined by Kaptive (29)), the middle ring the *kpsT* variant and the inner ring the *kpsM* variant. The divergent alleles fall in the magenta subclade identified as capsule type KL112.

There were 32 k-types, 14 known and 18 unknown and 32 KL types identified. The predominant ExPEC related capsule K5 – KL5 was found in 977 isolates, and the K100 – KL100 in 608 isolates. The K100 – KL100 capsule type dominated clade A and was also found in a sub-clade of clade C2 which contained the ST131 reference genome EC958. A phylogenetically distinct cluster of genomes was identified within clade C with an unknown atypical capsule type assigned KL112 (highlighted in Figure 2). This KL112 capsule type was not found in a wider search of our genome collection of 20,577 *E. coli* genomes from 20 other STs but present in public genomes outside our curated set. The 152 isolates containing the KL112 capsule type were identified between 2003 – 2016, predominantly from North America (n=41) and Europe (n=37) but also found in Asia (n=8), Africa (n=5) and Oceania (n=1). The source of the isolates was mostly human (n=99) including UTI, bacteraemia and asymptomatic. Further metadata was not available.

To fully characterise this unique capsule region, we conducted comparative long read genome sequencing on three isolates in our collection from this KL112 cluster and two others from the ST131 phylogeny: USVAST065 (KL115) from clade A and USVAST219 (KL5) from clade B. This comparison also included ST131 reference strain EC958 (KL100) which is located in clade C. Long read sequence analysis revealed an abnormal arrangement of the *kps* operon in the KL112 genomes (Fig 3).

**Figure 3.**
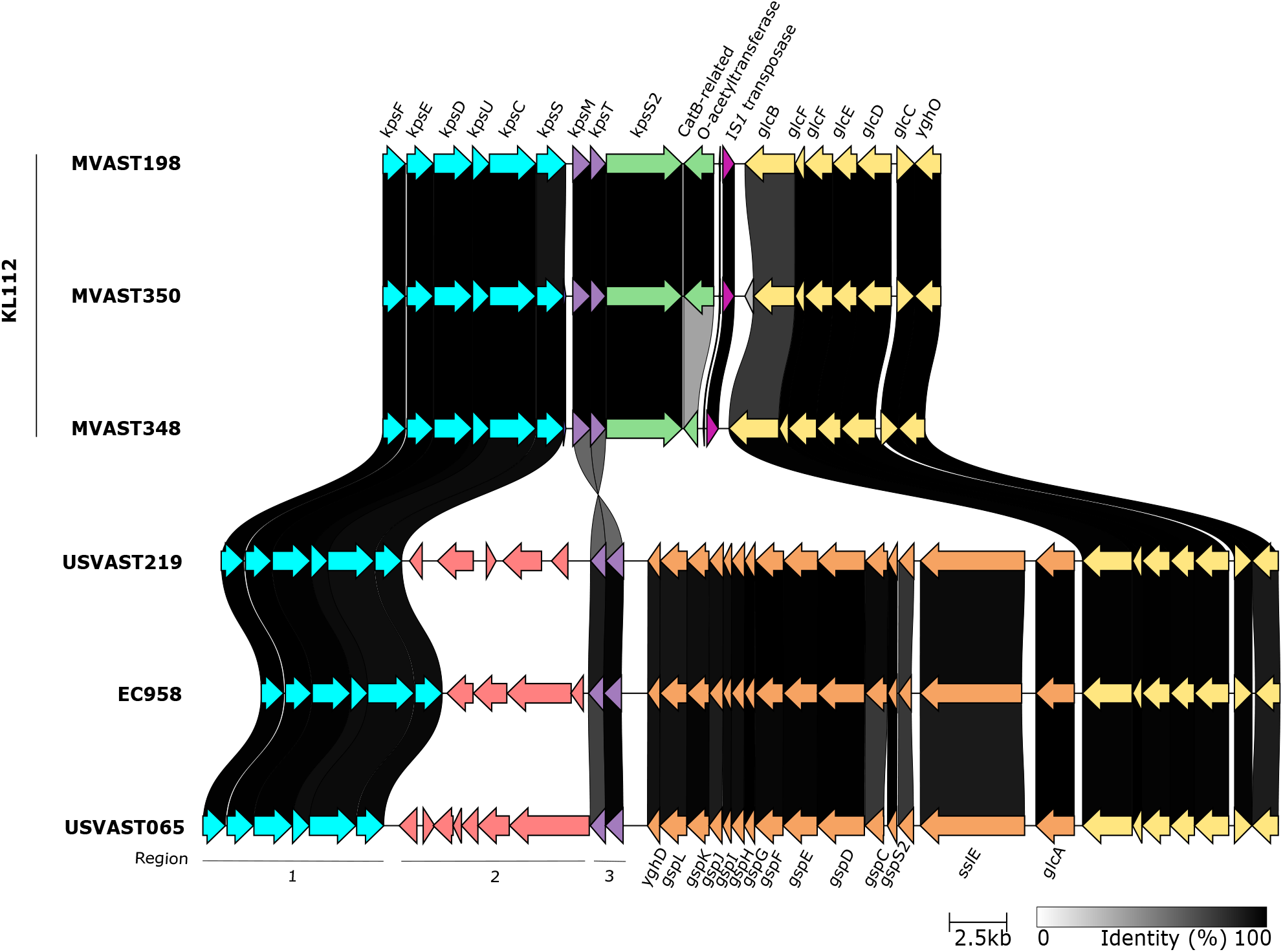
Genetic organisation of the *kps* operon of six ST131 *E. coli* isolates. The top three isolates (MVAST198 (33), MVAST350 (33) and MVAST348 (33)) contain the divergent *kpsM* allele and rearrangement which demonstrates a rotation of the *kpsM* and *kpsT* genes, as well as the deletion of the *gsp* operon at this locus. The bottom three isolates (USVAST219 (34), EC958 (25), USVAST065 (34)) contain the conventional arrangement.

In the genome of KL112 strains, the *kpsM* and *kpsT* genes are inverted such that they are in the same orientation as the other *kps* genes in the operon. The region 2 genes reside on the outside of the operon downstream of the reversed *kpsMT* pair. There has also been loss of the PR3 promoter and the ops originally upstream of the *kpsM* gene which normally allow for the transcription of *kpsM* and *kpsT* and extension to region 2 genes despite their differing orientation to the rest of the operon. *kpsS* is 105 bp longer in the KL112 type genomes. A copy of the 768 bp insertion sequence IS*1* is located downstream of *glcB*. This IS*1* is not flanked by a target site duplication, which is consistent with an IS*1*-mediated deletion event subsequent to its insertion here. Between *kpsMT* and IS*1* lies a second *kps*S gene (*kpsS*2) and an open reading frame for a putative O-acetyltransferase related to CatB. The *gsp* operon, normally found downstream of the *kps* operon, is not present adjacent to the capsule locus in the KL112 type, and might have been lost from this position in an IS*1*-mediated deletion event or as a result of substantial recombination.

### *kpsM* and *kpsT* rearrangement does not aGect transcription of the capsule region

RNAseq was performed on four isolates, three with the KL112 type abnormal arrangement and one with the conventional *kpsMT* arrangement to assess whether the *kpsM* and *kpsT* genes in the operon were still expressed. Read counts relative to housekeeping gene *adk* showed *kpsM* and *kpsT* were expressed in both arrangements (Fig 4). In both operon configurations, the *kpsM* and *kpsT* genes were expressed at lower levels than the Region 1 genes. The IS*1* in this region likely provides an outward facing promoter leading to the expression of *kpsS2* and the *catB*-related O-acetyltransferase.

**Figure 4.**
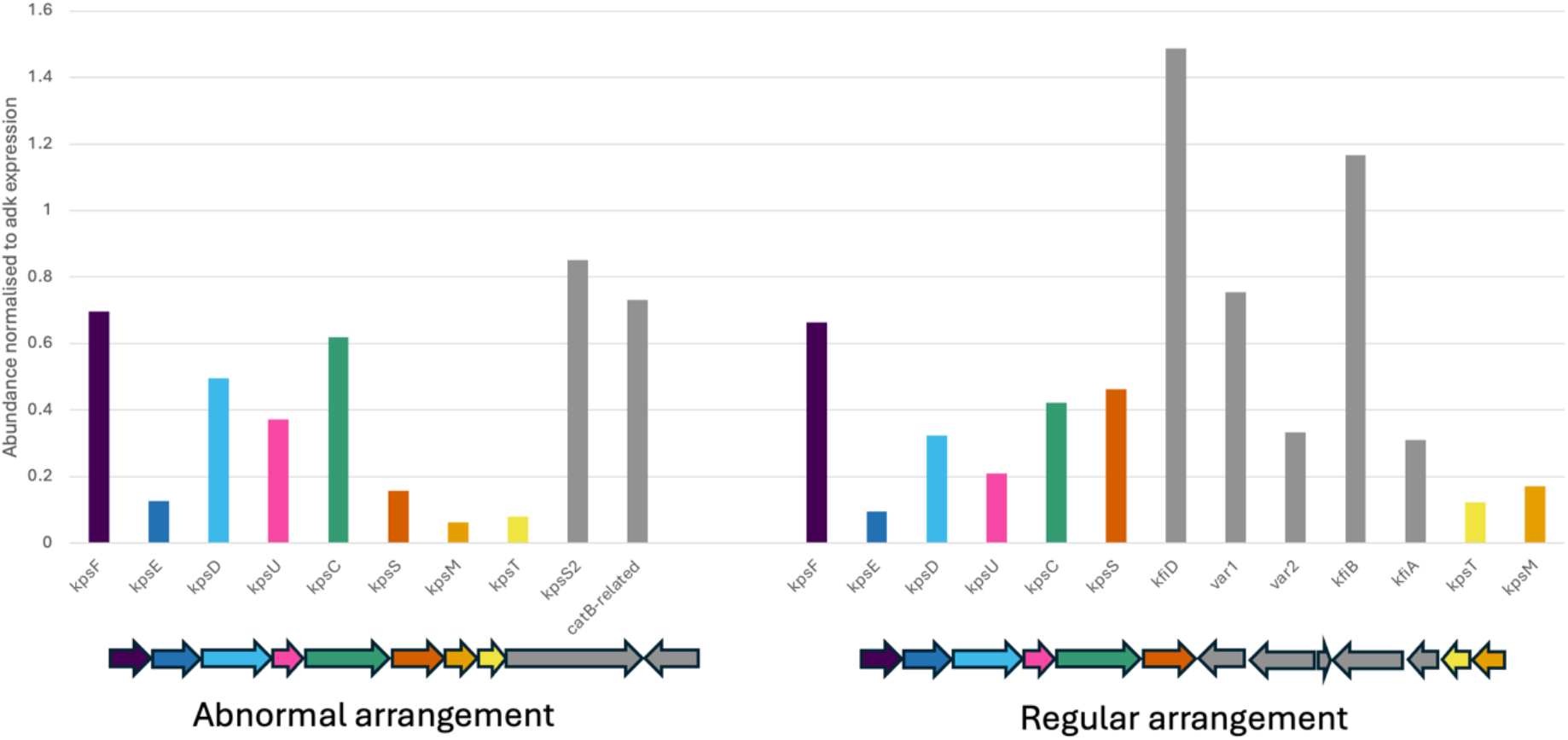
Gene expression of the *kps* operon in both the abnormal arrangement and the regular arrangement. RNAseq was carried out to identify if genes *kpsM* (orange) and *kpsT* (yellow) are still expressed in the rearrangement. Results were normalised to expression of housekeeping gene *adk*.

**Figure 5.**
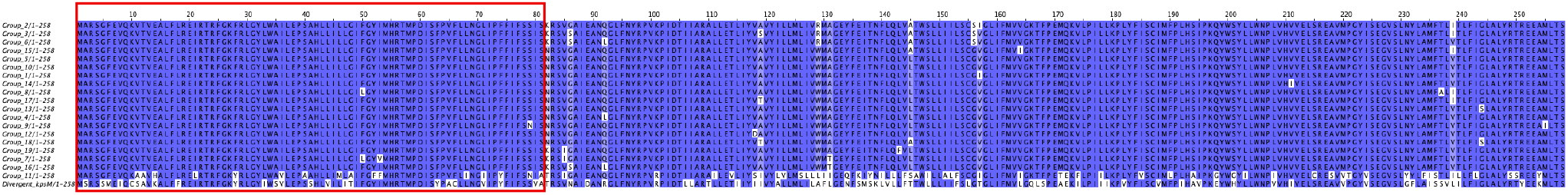
Amino acid sequence alignment of KpsM variants found in ST131. Variants were first clustered at a nucleotide level and a translated representative amino acid sequence was taken from each group. The red box is highlighting the relatively conserved first third of the protein.

### Sequence analysis of the KL112 *kpsMT* genes identifies alleles of these genes unique to the KL112 phylogenetic cluster of ST131

We extracted the sequences of *kpsM* and *kpsT* from our collection of 3,033 ST131 isolates. There were 52 different alleles of *kpsM* which we grouped into 21 clusters (supplementary figure 1), as described in Methods. A complete, annotated *kpsM* gene was detected in 2,912 genomes, 75% of these genomes (n = 2,182) grouped into one of the three *kpsM* clusters (Group 1, Group 2 and Group 3). A complete and annotated *kpsT* gene was present in 2,839 genomes consisting of 58 *kpsT* alleles, with 75% of genomes (n = 2,136) grouping into one of three *kpsT* allele clusters (Group 1, Group 2 and Group 3). These clusters correlate between *kpsM* and *kpsT*, with Group 1/Group 1, Group 2/Group 2, and Group 3/Group 3 allele clusters being found together in isolates (Fig 2) with an adjusted rand index of 0.954.

The *kpsM* alleles were all the same length at 777bp; apart from in one isolate where an early stop codon reduced the coding region to 474 bp. The *kpsT* alleles however, varied in length ranging from 600 bp to 708 bp. The gene length reported in GenBank is 676 bp. The KL112 allele of *kpsM* was the most divergent from the EC958 (29) reference genome allele, with just 62.9% nucleotide identity and an amino acid sequence identity of 80%. When comparing this allele to the other variants in ST131 the SNPs were distributed throughout the gene. For 50 of the variants, the first 240bp of the alignment is conserved with 28 sites (0.1%) showing variation with the latter half of the sequence showing variation in 106 sites over 537bp (0.2%). In the predicted structure of the KpsM protein, the conserved 240 bp of *kpsM* codes for the N-terminal tail, transmembrane helix 1, periplasmic loop 1 and transmembrane helix 2 (43).

Performing BLAST searches with the KL112 *kpsM* allele returned just 25 matches found in publicly available *E. coli* genomes, belonging to STs including phylogroup B2 ST131 (n=20), ST104 (n=1) and ST12285 (n=1) and phylogroup D ST6131 (n=2) and ST4353 (n=1). Performing the same search but excluding *E. coli* rendered no significant results. The alternative arrangement of the *kpsM* and *kpsT* genes can be found in other atypical K-loci identified in Kaptive but no other atypical K-loci were found in our ST131 collection. Comparing capsule type to the variants of *kpsM* and *kpsT* in each isolate showed that the variant type was mostly concordant with capsule type (Fig 2) with *kpsM* having an adjusted rand index of 0.829 with KL/K-type and *kpsT* an adjusted rand index of 0.906 with KL/K-type. This increased to an adjusted rand index of 0.922 and 0.939 when comparing the K/KL-type to the clustered *kpsM* groups and *kpsT* groups respectively. The exceptions came in eight of the 32 KL types identified. Seven of these showed a dominant group as well as a couple of variants for other groups. KL121 however, showed a spread of five different *kpsM* and *kpsT* variant groups. However, this locus’ match type is unknown suggesting uncertainty about its conformation.

## Discussion

In this study we reveal a distinctive clone within ST131 clade C. This clone is characterised by an inversion event in the region 3 capsule genes *kpsM* and *kpsT* which is basal to the monophyletic clone. This inversion results in a capsule-type switch within the clone. Clade C is estimated to have emerged in the 1980s and is characterised by the switch in *fim* allele as well as specific *gyrA* and *parC* mutations conferring resistance to fluoroquinolones (20). These evolutionary changes contribute to the proliferation of clade C with our study suggesting there is continued ongoing evolution within clade C shaping host-pathogen interaction, as similarly seen in *Klebsiella pneumoniae* (44).

*E. coli* is known to have a large diversity of capsule types (6, 7) with capsule switch events facilitated by the locus being part of a recombination hotspot (19, 45). A further typing scheming has been released – kTYPr ( https://github.com/SushiLab/kTYPr) (7). We have included the results of this in the supplementary material. Results were largely concordant to Gladstone *et al*. albeit with different naming scheme. By analysing thousands of ST131 genomes we show that although there is a large amount of gene gain and rearrangement in the capsule region of the genome, successful capsule types are selected and maintained, resulting in a clonal distribution across the lineage phylogeny. The flexibility within the capsule locus of ST131 might have aided the lineage’s success (6), as seen across other bacteria. Capsule switching has been well documented within several species as an effective measure for serotype replacement which can lead to vaccine avoidance, as well as immune evasion. Following the introduction of the pneumococcal conjugate vaccine in 2003, *Streptococcus pneumoniae* serotypes not protected against in the vaccine, emerged in STs they had never been seen in before, despite extensive surveillance (46, 47). Genomic investigation revealed a recombination event at the capsular locus. This capsule switch provided these STs significant evolutionary advantages and led to the increase in prevalence of capsule-switched strains (48). Similarly, in *Neisseria meningitidis* there is evidence of increased immune evasion and vaccine avoidance by capsule switching (49, 50).

Our findings suggest that an ancestral capsule switch event has defined this clone within ST131 clade C. The capsule type is isolated to this single monophyletic region of the ST131 phylogenetic tree with a few one-off occurrences in other less common *E. coli* lineages. The same applies to the divergent *kpsMT* alleles within this capsule type. Furthermore, our investigation could find no other notable genetic feature to define this clone, such as acquisition of resistance genes or O-antigen switching. There are a number of similar ancestral events that have defined the formation of dominant clones in *E. coli* lineages across the species. Examples of this include the O-antigen switch in ST167 as well as the O-antigen switch and recombination of novel iron acquisition alleles in ST410 (51). ST167 emerged from clonal complex ST10 with a notable loss of genes controlling the length of the O-antigen, resulting in a truncated O-antigen (52). Clinical cases of these bacteria are found across the globe, frequently carrying both Extended Spectrum Beta-Lactamase and carbapenemase-producing genes (52). In ST410, a hypervirulent MDR clone emerged with increasing cases worldwide that was noted to have switched to a novel O-antigen and to have gained a high pathogenicity island that enhances iron acquisition (53), allowing for adaptation to changing environments. Therefore, it is essential to monitor such events in bacterial populations as it may allow us to capture the formation of successful clones.

Capsule typing focusses on the region 2 genes of the capsule operon that vary between capsule types, however our research also suggests the *kpsM* and *kpsT* gene alleles are capsule specific as *kpsM* and *kpsT* alleles appear concordant with capsule type across the ST131 phylogenetic tree. *kpsM* is the integral inner membrane component and *kpsT* the ATPase, which together form the ATP-binding cassette that transports the capsular polysaccharides across the inner membrane. The concordance of *kpsM* and *kpsT* alleles with capsule types could suggest the different variants affect the transportation of different polysaccharides, and therefore the change in capsule phenotype.

Our results are suggestive that the event leading to the atypical configuration of the capsule operon and the divergent *kpsM* and *kpsT* alleles occurred *de novo* in this ST131 clone with rare recombination events resulting in its presence in other STs. This clone does not appear to have become dominant in humans. The likelihood is that this ancestral capsule switching event led to improved immune or phage evasion for this specific clone, but that a wider selective fitness advantage is not conferred leading to its relative confinement in this specific monophyletic clade of ST131.

## Acknowledgements

This work was supported by the Biotechnology and Biological Sciences Research Council (BBSRC) and University of Birmingham funded Midlands Integrative Biosciences Training Partnership (MIBTP) (grant number BB/T00746X/1) through a PhD studentship awarded to IP. RJH and EAC are funded by a BBSRC grant (BB/W020602/1) awarded to AM. RAM was funded by NERC grant (NE/T01301X/1) awarded to AM. AM is funded by the National Institute for Health and Care Research (NIHR) Birmingham Biomedical Research Centre – NIHR203326.

## Data summary

The genome sequence data generated in this study are publicly available from NCBI under BioProject PRJNA1289411 and PRJNA1330052. The *Escherichia coli* genome data sets used in this work are from a previous publication (23), ENA accessions and supporting metadata can be found in the supplementary table. The RNAseq data is available from NCBI under BioProject PRJNA1369585.

## Impact statement

Multi-drug resistant (MDR) Escherichia coli is a problematic cause of invasive disease in humans and animals. Resistance to multiple classes of antibiotics is severely limiting treatment options. ST131-C2 is a highly successful, MDR clone of *E. coli* that is a major cause of extraintestinal infections. Capsule is a key virulence factor in contributing to this clones success with capsule switching allowing the bacteria flexibility to thrive in different niches. We present evidence that suggests continual evolution of this proliferant clone influenced by host (44)-pathogen interaction. Small genetic changes can define the success of a clone, therefore it is important to monitor this continual evolution in pathogens.

## References

1. Whitfield C. Biosynthesis and assembly of capsular polysaccharides in Escherichia coli. Annu Rev Biochem. 2006;75:39–68.

2. Whitfield C, Wear SS, Sande C. Assembly of Bacterial Capsular Polysaccharides and Exopolysaccharides. Annu Rev Microbiol. 2020;74:521–43.

3. Kuklewicz J, Zimmer J. Molecular insights into capsular polysaccharide secretion. Nature. 2024;628(8009):901–9.

4. Cress BF, Englaender JA, He W, Kasper D, Linhardt RJ, Koffas MA. Masquerading microbial pathogens: capsular polysaccharides mimic host-tissue molecules. FEMS Microbiol Rev. 2014;38(4):660–97.

5. Sande C, Bouwman C, Kell E, Nickerson NN, Kapadia SB, Whitfield C. Structural and Functional Variation in Outer Membrane Polysaccharide Export (OPX) Proteins from the Two Major Capsule Assembly Pathways Present in Escherichia coli. J Bacteriol. 2019;201(14).

6. Gladstone RA, Pesonen M, Pontinen AK, Maklin T, MacAlasdair N, Thorpe H, et al. Identification of transporter-dependent capsular loci associated with the invasive potential of Escherichia coli. Nat Microbiol. 2026.

7. Miravet-Verde S, Cacace E, Mores CR, Rutschmann C, Lin CW, Ruscheweyh HJ, et al. In silico typing maps the natural diversity of Escherichia coli transporter-dependent capsules. Nat Microbiol. 2026.

8. Clarke BR, Pearce R, Roberts IS. Genetic organization of the Escherichia coli K10 capsule gene cluster: identification and characterization of two conserved regions in group III capsule gene clusters encoding polysaccharide transport functions. J Bacteriol. 1999;181(7):2279–85.

9. Whitney JC, Howell PL. Synthase-dependent exopolysaccharide secretion in Gram-negative bacteria. Trends Microbiol. 2013;21(2):63–72.

10. Sande C, Whitfield C. Capsules and Extracellular Polysaccharides in Escherichia coli and Salmonella. EcoSal Plus. 2021;9(2):eESP00332020.

11. Jann K, Jann B. Capsules of Escherichia coli, expression and biological significance. Can J Microbiol. 1992;38(7):705–10.

12. Korhonen TK, Valtonen MV, Parkkinen J, Vaisanen-Rhen V, Finne J, Orskov F, et al. Serotypes, hemolysin production, and receptor recognition of Escherichia coli strains associated with neonatal sepsis and meningitis. Infect Immun. 1985;48(2):486–91.

13. Whitfield C, Roberts IS. Structure, assembly and regulation of expression of capsules in Escherichia coli. Mol Microbiol. 1999;31(5):1307–19.

14. Willis LM, Whitfield C. KpsC and KpsS are retaining 3-deoxy-D-manno-oct-2-ulosonic acid (Kdo) transferases involved in synthesis of bacterial capsules. Proc Natl Acad Sci U S A. 2013;110(51):20753–8.

15. Bliss JM, Garon CF, Silver RP. Polysialic acid export in Escherichia coli K1: the role of KpsT, the ATP-binding component of an ABC transporter, in chain translocation. Glycobiology. 1996;6(4):445–52.

16. Xue P, Corbett D, Goldrick M, Naylor C, Roberts IS. Regulation of expression of the region 3 promoter of the Escherichia coli K5 capsule gene cluster involves H-NS, SlyA, and a large 5’ untranslated region. J Bacteriol. 2009;191(6):1838–46.

17. Jia J, King JE, Goldrick MC, Aldawood E, Roberts IS. Three tandem promoters, together with IHF, regulate growth phase dependent expression of the Escherichia coli kps capsule gene cluster. Sci Rep. 2017;7(1):17924.

18. Aldawood E, Roberts IS. Regulation of Escherichia coli Group 2 Capsule Gene Expression: A Mini Review and Update. Front Microbiol. 2022;13:858767.

19. Alqasim A, Scheutz F, Zong Z, McNally A. Comparative genome analysis identifies few traits unique to the Escherichia coli ST131 H30Rx clade and extensive mosaicism at the capsule locus. BMC Genomics. 2014;15(1):830.

20. Petty NK, Ben Zakour NL, Stanton-Cook M, Skippington E, Totsika M, Forde BM, et al. Global dissemination of a multidrug resistant Escherichia coli clone. Proc Natl Acad Sci U S A. 2014;111(15):5694–9.

21. Goh KGK, Phan MD, Forde BM, Chong TM, Yin WF, Chan KG, et al. Genome-Wide Discovery of Genes Required for Capsule Production by Uropathogenic Escherichia coli. mBio. 2017;8(5).

22. Zhou Z, Alikhan NF, Mohamed K, Fan Y, Agama Study G, Achtman M. The EnteroBase user’s guide, with case studies on Salmonella transmissions, Yersinia pestis phylogeny, and Escherichia core genomic diversity. Genome Res. 2020;30(1):138–52.

23. Cummins EA, Hall RJ, Connor C, McInerney JO, McNally A. Distinct evolutionary trajectories in the Escherichia coli pangenome occur within sequence types. Microb Genom. 2022;8(11).

24. Altschul SF, Gish W, Miller W, Myers EW, Lipman DJ. Basic local alignment search tool. J Mol Biol. 1990;215(3):403–10.

25. Camacho C, Coulouris G, Avagyan V, Ma N, Papadopoulos J, Bealer K, et al. BLAST+: architecture and applications. BMC Bioinformatics. 2009;10:421.

26. Stothard P. The sequence manipulation suite: JavaScript programs for analyzing and formatting protein and DNA sequences. Biotechniques. 2000;28(6):1102, 4.

27. Katoh K, Rozewicki J, Yamada KD. MAFFT online service: multiple sequence alignment, interactive sequence choice and visualization. Brief Bioinform. 2019;20(4):1160–6.

28. Waterhouse AM, Procter JB, Martin DM, Clamp M, Barton GJ. Jalview Version 2--a multiple sequence alignment editor and analysis workbench. Bioinformatics. 2009;25(9):1189–91.

29. Forde BM, Ben Zakour NL, Stanton-Cook M, Phan MD, Totsika M, Peters KM, et al. The complete genome sequence of Escherichia coli EC958: a high quality reference sequence for the globally disseminated multidrug resistant E. coli O25b:H4-ST131 clone. PLoS One. 2014;9(8):e104400.

30. Madeira F, Madhusoodanan N, Lee J, Eusebi A, Niewielska A, Tivey ARN, et al. The EMBL-EBI Job Dispatcher sequence analysis tools framework in 2024. Nucleic Acids Res. 2024;52(W1):W521–W5.

31. Jolley KA, Maiden MC. BIGSdb: Scalable analysis of bacterial genome variation at the population level. BMC Bioinformatics. 2010;11:595.

32. Beghain J, Bridier-Nahmias A, Le Nagard H, Denamur E, Clermont O. ClermonTyping: an easy-to-use and accurate in silico method for Escherichia genus strain phylotyping. Microb Genom. 2018;4(7).

33. Lam MMC, Wick RR, Judd LM, Holt KE, Wyres KL. Kaptive 2.0: updated capsule and lipopolysaccharide locus typing for the Klebsiella pneumoniae species complex. Microb Genom. 2022;8(3).

34. Tonkin-Hill G, MacAlasdair N, Ruis C, Weimann A, Horesh G, Lees JA, et al. Producing polished prokaryotic pangenomes with the Panaroo pipeline. Genome Biol. 2020;21(1):180.

35. Stamatakis A. RAxML version 8: a tool for phylogenetic analysis and post-analysis of large phylogenies. Bioinformatics. 2014;30(9):1312–3.

36. Letunic I, Bork P. Interactive Tree Of Life (iTOL): an online tool for phylogenetic tree display and annotation. Bioinformatics. 2007;23(1):127–8.

37. Scrucca L, Fop M, Murphy TB, Raftery AE. mclust 5: Clustering, Classification and Density Estimation Using Gaussian Finite Mixture Models. R J. 2016;8(1):289–317.

38. Biggel M, Xavier BB, Johnson JR, Nielsen KL, Frimodt-Moller N, Matheeussen V, et al. Horizontally acquired papGII-containing pathogenicity islands underlie the emergence of invasive uropathogenic Escherichia coli lineages. Nat Commun. 2020;11(1):5968.

39. Colpan A, Johnston B, Porter S, Clabots C, Anway R, Thao L, et al. Escherichia coli sequence type 131 (ST131) subclone H30 as an emergent multidrug-resistant pathogen among US veterans. Clin Infect Dis. 2013;57(9):1256–65.

40. Bouras G, Houtak G, Wick RR, Mallawaarachchi V, Roach MJ, Papudeshi B, et al. Hybracter: enabling scalable, automated, complete and accurate bacterial genome assemblies. Microb Genom. 2024;10(5).

41. Schwengers O, Jelonek L, Dieckmann MA, Beyvers S, Blom J, Goesmann A. Bakta: rapid and standardized annotation of bacterial genomes via alignment-free sequence identification. Microb Genom. 2021;7(11).

42. Carver T, Harris SR, Berriman M, Parkhill J, McQuillan JA. Artemis: an integrated platform for visualization and analysis of high-throughput sequence-based experimental data. Bioinformatics. 2012;28(4):464–9.

43. Pigeon RP, Silver RP. Topological and mutational analysis of KpsM, the hydrophobic component of the ABC-transporter involved in the export of polysialic acid in Escherichia coli K1. Mol Microbiol. 1994;14(5):871–81.

44. Le Bris J, Varet H, Rocha EPC, Rendueles O. Plug-and-play evolution of the Klebsiella pneumoniae capsule locus enables serotype exchange across genetic backgrounds. PLoS Biol. 2026;24(3):e3003724.

45. Touchon M, Hoede C, Tenaillon O, Barbe V, Baeriswyl S, Bidet P, et al. Organised genome dynamics in the Escherichia coli species results in highly diverse adaptive paths. PLoS Genet. 2009;5(1):e1000344.

46. Brueggemann AB, Pai R, Crook DW, Beall B. Vaccine escape recombinants emerge after pneumococcal vaccination in the United States. PLoS Pathog. 2007;3(11):e168.

47. Croucher NJ, Finkelstein JA, Pelton SI, Mitchell PK, Lee GM, Parkhill J, et al. Population genomics of post-vaccine changes in pneumococcal epidemiology. Nat Genet. 2013;45(6):656–63.

48. Sabharwal V, Stevenson A, Figueira M, Orthopoulos G, Trzcinski K, Pelton SI. Capsular switching as a strategy to increase pneumococcal virulence in experimental otitis media model. Microbes Infect. 2014;16(4):292–9.

49. Swartley JS, Marfin AA, Edupuganti S, Liu LJ, Cieslak P, Perkins B, et al. Capsule switching of Neisseria meningitidis. Proc Natl Acad Sci U S A. 1997;94(1):271–6.

50. Beddek AJ, Li MS, Kroll JS, Jordan TW, Martin DR. Evidence for capsule switching between carried and disease-causing Neisseria meningitidis strains. Infect Immun. 2009;77(7):2989–94.

51. Feng Y, Liu L, Lin J, Ma K, Long H, Wei L, et al. Key evolutionary events in the emergence of a globally disseminated, carbapenem resistant clone in the Escherichia coli ST410 lineage. Commun Biol. 2019;2:322.

52. Zong Z, Fenn S, Connor C, Feng Y, McNally A. Complete genomic characterization of two Escherichia coli lineages responsible for a cluster of carbapenem-resistant infections in a Chinese hospital. J Antimicrob Chemother. 2018;73(9):2340–6.

53. Ba X, Guo Y, Moran RA, Doughty EL, Liu B, Yao L, et al. Global emergence of a hypervirulent carbapenem-resistant Escherichia coli ST410 clone. Nat Commun. 2024;15(1):494.

