## Supplementary Figures for "Genomic rearrangement of the capsule operon in a phylogenetically distinct cluster of the multi-drug resistant *Escherichia coli* lineage ST131"

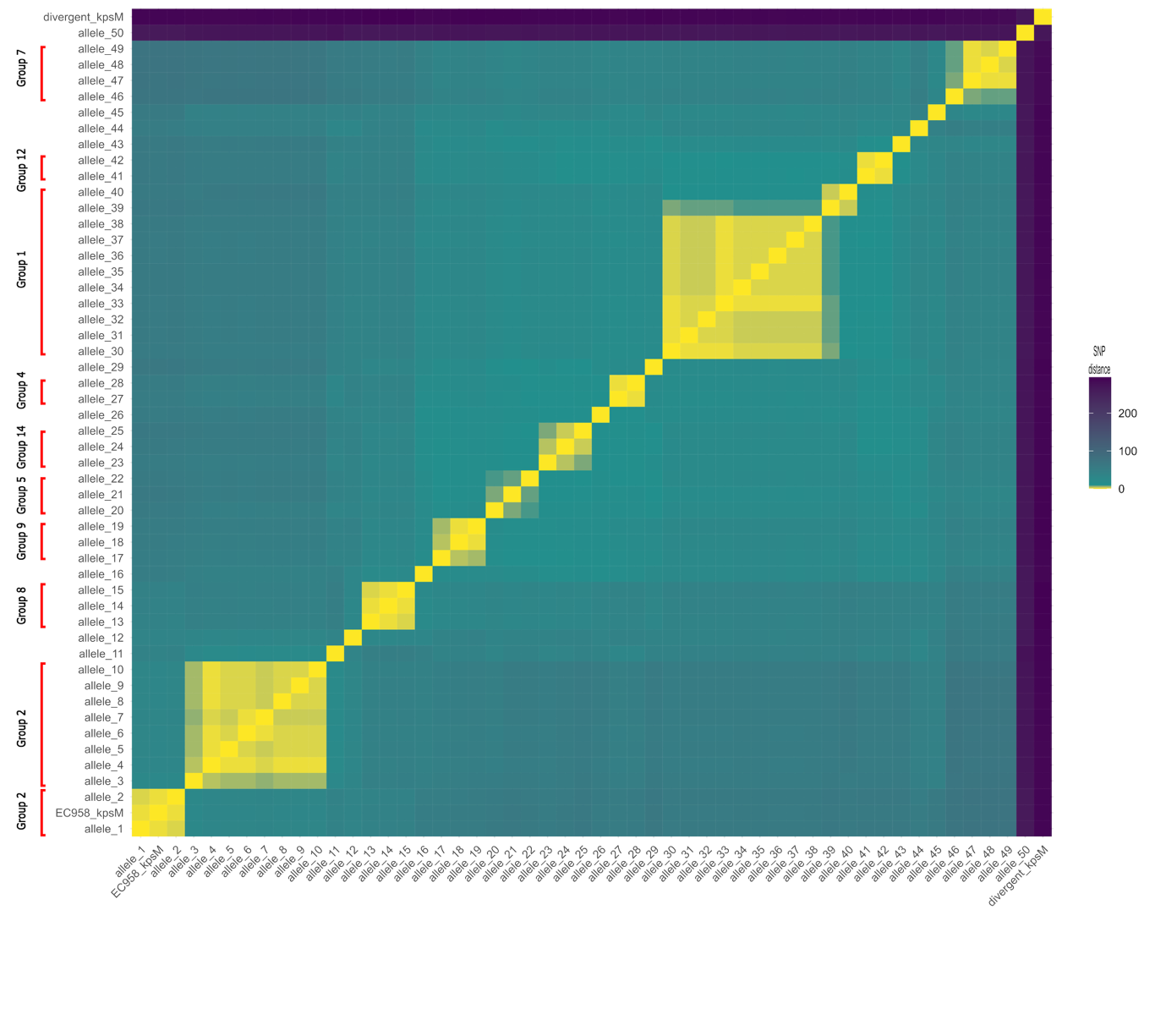

**Supplementary figure 1** A heatmap showing the SNP distance calculated with snp-dists (v0.8.2) (<https://github.com/tseemann/snp-dists>) between kpsM alleles identified in a collection of 3,033 ST131 E. coli. These alleles were then clustered into groups according to their SNP distance.
